# Visualization of peripheral nerves in developing and regenerating limbs using a novel peripherin reporter line of *Xenopus laevis*

**DOI:** 10.64898/2026.04.19.719517

**Authors:** Miyuki Suzuki, Yosuke Kato, Rima Mizuno, Hiroshi Yajima, Shinichirou Miura, Tetsuya Endo, Makoto Mochii, Ken-ichi T. Suzuki

**Affiliations:** Division of Biology and Biological Engineering, California Institute of Technology, California, 91125, United States; Department of Life Science, Graduate School of Science, University of Hyogo, Akougun, Hyogo, 678-1297, Japan; Department of Basic Biology, The Graduate University for Advanced Studies, SOKENDAI, Okazaki, Aichi, 444-8585, Japan; Gotemba Nishi High School,644-1 Gumisawa, Gotemba, Shizuoka 412-0041, Japan; Division of Liberal Arts and Sciences, Aichi Gakuin University, Nissin, Aichi, 470-0195, Japan; Trans-Scale Biology Center, National Institute for Basic Biology, Okazaki, Aichi, 444-8585, Japan

**Keywords:** limb, peripherin, nerve, metamorphosis, regeneration, transgenic, *Xenopus laevis*

## Abstract

Peripherin (*PRPH*) is a class III intermediate filament protein expressed in peripheral nerves and upregulated during axon outgrowth and regeneration. In this study, we developed a transgenic *Xenopus laevis* line for long-term *in vivo* visualization of the peripheral nervous system. Deletion and motif analyses identified *cis*-regulatory regions within the promoter and intron 1 that are important for neuronal expression of the *X. laevis prph* gene. Stable lines exhibited robust EGFP reporter activity in developing neural primordia in embryos and in the peripheral nerves of tadpoles. Transgenic tadpoles enabled *in vivo* imaging of peripheral nerves throughout limb development. During larval limb regeneration, we observed modest early nerve entry into the blastema, recapitulating that seen in early limb development. In contrast, post-metamorphic limb blastemas displayed extensive innervation from the early phase of regeneration. Moreover, increased reporter activity in the nerves of the regenerating adult forelimb suggests regeneration-associated regulation of peripheral innervation and its potential role in blastema formation. This transgenic line will serve as a versatile tool for analyzing such large-scale neural remodeling across development, metamorphosis, and regeneration.

## 1. Introduction

Peripherin (*PRPH*) is a class III intermediate filament protein (Portier et al., 1983) that is expressed in the peripheral nervous system (PNS), particularly in sensory and autonomic neurons (Brody et al., 1989; Escurat et al., 1990; Yuan et al., 2012). *PRPH* is highly conserved among vertebrates, including mammals, birds, reptiles, amphibians, and fish. In *Xenopus laevis*, the *prph* gene (also known as XIF3) was reported to be expressed in a broad range of PNS and central nervous system (CNS) structures, including the neural plate primordium, olfactory placodes, cranial nerve ganglia in the neurula stage embryo, and the brain and spinal cord at tailbud stage (Green and Vetter, 2011; Sharpe et al., 1989). Rat *Prph* and human *PRPH* expression are known to increase during the outgrowth of axons in neural development (Escurat et al., 1990; Xiao et al., 2006), suggesting an important role in axon growth. In rat PC12 cells, the expression of *Prph* specifically increased during nerve growth factor (NGF)-promoted neuronal differentiation (Aletta et al., 1989; Leonard et al., 1988). In dissociated embryonic spinal cord cultures of *X. laevis*, Prph is actively expressed in immature neurite branches (Smith et al., 2006; Undamatla and Szaro, 2001). *PRPH* upregulation is also well recognized during axon regeneration, where damaged peripheral neurons show a substantial increase in its expression. This expression is subsequently downregulated once elongation is complete, supporting its potential role in axon regrowth (Oblinger et al., 1989; C.M. Troy et al., 1990; Wong and Oblinger, 1990). In addition, *PRPH* has been associated with polyneuropathies, traumatic nerve injury, and diabetic neuropathy, and has recently gained attention as a potential biomarker for nerve damage in the PNS (Hol and Capetanaki, 2017; Keddie et al., 2023; Manco et al., 2025; Romano et al., 2022).

Anuran amphibians including *X. laevis* can regenerate a variety of organs, such as limbs, tail, and parts of the brain, both morphologically and functionally in larvae (Beck et al., 2009; Dent, 1962; Yoshino and Tochinai, 2006). However, *X. laevis* undergoes robust tissue remodeling during metamorphosis from larva to adult and loses the regeneration capability. While tadpoles are capable of both scar-free wound healing and tissue regeneration, these capabilities are impaired in adulthood (Phipps et al., 2020). In particular, tadpoles can regenerate hindlimbs during the pre-metamorphic stage (st. 50–52), but the number of regenerated digits decreases in later stages, and post-metamorphic froglets form only a hypomorphic cartilage spike (Dent, 1962; Endo et al., 2004; Keenan and Beck, 2016; Tassava, 2004). In urodele amphibians (salamanders), which exhibit remarkable regeneration capabilities even in adulthood, nerves play a critical role in limb regeneration; denervation results in total regenerative failure, and it has been suggested that nerve-derived trophic factors support blastema cell survival and proliferation (Kumar and Brockes, 2012; Pirotte et al., 2016; Stocum, 2017). Therefore, a detailed analysis of innervation is necessary for understanding limb regeneration in *Xenopus* as well.

For continuous *in vivo* analysis of post-embryonic processes, including metamorphosis and regeneration, transgenic *Xenopus* lines visualizing nerves are poised to become an essential and versatile resource for PNS and organ regeneration studies. For visualizing neurons, transgenic frogs expressing reporters under the control of the *β-tubulin class II* (*tubb2b*) promoter have been used and cover nearly all neuronal populations in the CNS (Horb et al., 2019; Love et al., 2011). In the *tubb2b*::GFP transgenic *Xenopus* lines, numerous GFP-positive neurons are observed to be distributed in the regenerating blastema of froglets (Suzuki et al., 2006). However, it was reported that a subset of neurons was not labeled in the forebrain of *tubb2b* transgenic line (Daume et al., 2022), suggesting the benefits of expanding the range of neuronal reporters focusing on the PNS, in particular, dorsal root ganglion (DRG) derived from neural crest.

In this study, we aimed to establish a transgenic *X. laevis* line for *in vivo* imaging of the PNS, in particular, DRG neurons. Through sequential deletion analysis of the enhancer and promoter using EGFP reporter constructs, we narrowed down the essential region for neuron-specific transcription of the *prph* gene. We then established a stable transgenic line in which EGFP is driven by the *prph* promoter and intron 1, showing strong reporter expression in the neural plate during embryogenesis and in the PNS during larval stages. Using this line, we imaged peripheral nerve patterns during limb development and regeneration in both pre- and post-metamorphic stages, revealing distinct patterns of innervation between larval and adult stages. Thus, this study demonstrates *prph* as a valuable marker for *in vivo* studies of the PNS, especially for investigating nerve regeneration and metamorphosis.

## 2. Results

### 2.1. Establishment of *prph*:EGFP transgenic lines

Previous studies in mice demonstrated that intron 1 is essential for cell-type specific expression of rat *Prph* while it was not responsive to injury cues (Uveges et al., 2002). Comparative sequence analysis of the 4.7 kb intron 1 between the *X. laevis* (*prph*.*L*) and *X. tropicalis prph* genes using a VISTA plot identified four highly conserved regions (Fig. 1A). Similar conserved regions were also found in the *prph*.*S* (data not shown). Accordingly, we isolated the full-length intron 1 and a 914 bp promoter region including the 5’ UTR sequence of *prph*.*L* from a genomic library, and constructed a transgenic vector (Fig. 1B). Transgenic EGFP (*prph*:EGFP) fluorescence was first detected in the neural primordium, olfactory placode, and cranial ganglia at the neurula stage (Fig. 1C). At the tailbud stage, strong fluorescence was observed in the CNS, including the brain and spinal cord. Whole-mount *in situ* hybridization (WISH) detected *prph* mRNA expression in the embryonic regions corresponding to the EGFP reporter pattern, which is consistent with previous reports (Green and Vetter, 2011; Sharpe et al., 1989). In transgenic tadpoles, robust EGFP fluorescence was observed not only in CNS-but also in PNS-derived structures (e.g., optic nerve and trigeminal nerve; Fig. 1D).

**Figure 1.**
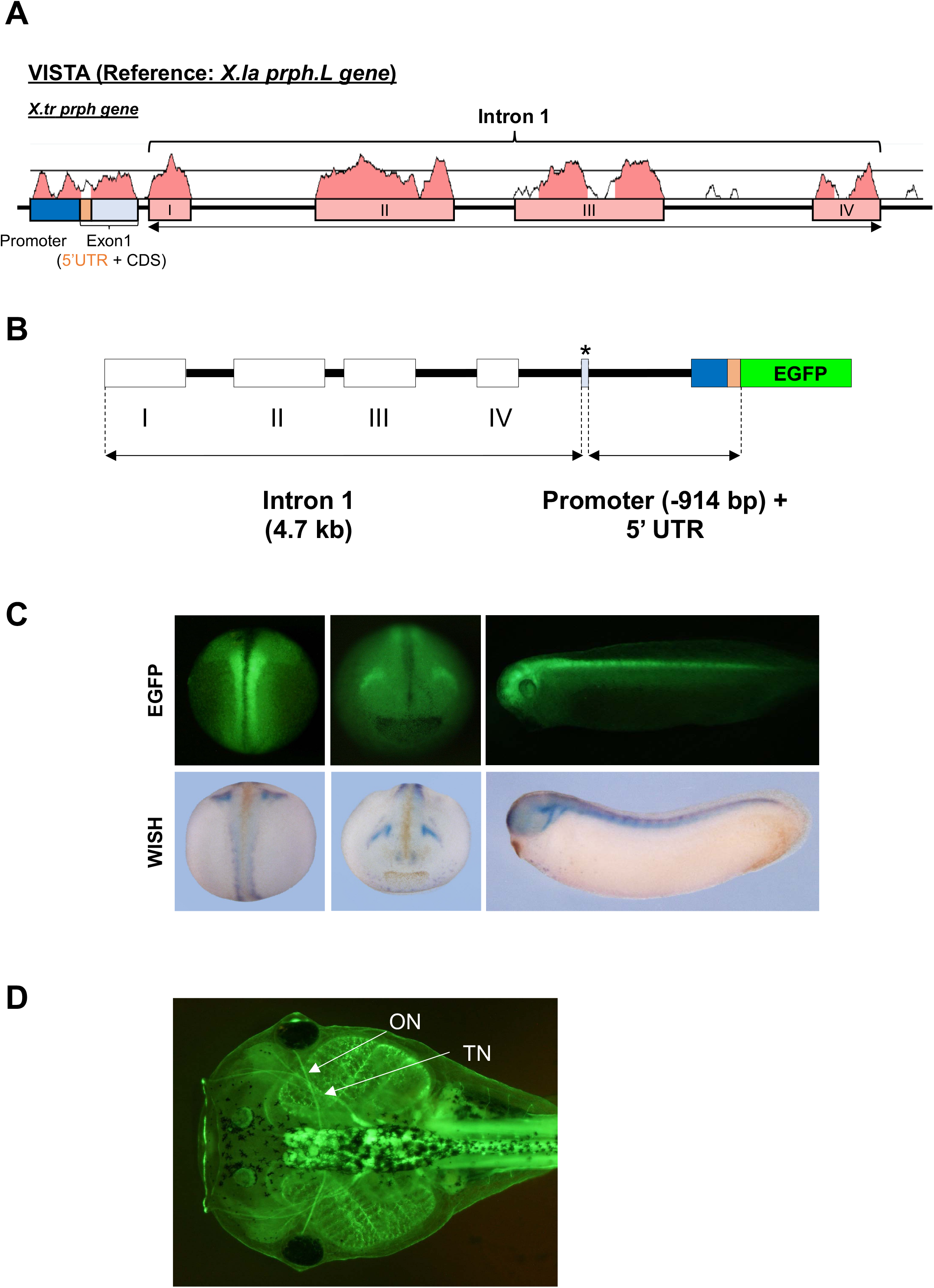
Expression of *prph*:EGFP and endogenous *prph* in embryos. (A) Comparative sequence analysis using VISTA plots. Sequence comparison of the *prph* gene between *X. laevis* (L subgenome) and *X. tropicalis* revealed conserved regions in the promoter and intron 1. (B) Structure of the construct used for transgenesis. White boxes indicate four regions of sequence homology between *Xenopus* species in intron 1. The *prph*.L promoter region containing exon 1 (914 bp, upstream region and 5’ UTR, blue and orange box, respectively), exon 2 (34 bp, a partial CDS, light blue box with an asterisk), and intron 1 (4.7 kb) was linked to EGFP. (C) Representative images of *prph*:EGFP transgenic embryos and whole-mount *in situ* hybridization (WISH) for endogenous *prph* expression. Neural cell-specific EGFP expression was observed in the neurula and tailbud stages, consistent with the *prph* expression pattern detected by WISH. (D) A representative reporter image of *prph*:EGFP transgenic *X. laevis* larva. Abbreviations: ON, optic nerve; TN, trigeminal nerve.

### 2.2. Sequential deletion analysis of the *prph*.*L* intron 1 enhancer

To further characterize the intron 1 enhancer, which consists of four conserved regions among *Xenopus prph* genes, we performed a sequential deletion analysis of the enhancer (Fig. 2A). Combining the full-length intron 1 with the CMV minimal promoter exhibited specific and strong EGFP fluorescence in the CNS at the tailbud stage. Deletion analysis was performed by removing each conserved region, which revealed that region II is essential for neural cell-specific enhancer activity in transgenic embryos. All deletion constructs tested are summarized in Fig. S1. To investigate the regulatory mechanism of *prph* activation in the peripheral neurons, we analyzed the chromatin accessibility of the *X. tropicalis prph* gene using the public ATAC-seq data at the neurula stage from Xenbase (Fisher et al., 2023). ATAC-seq peaks indicated chromatin accessibility at the promoter region and homology region II of intron 1 (Fig. 2B) at the neurula stage when *prph* is expressed. Motif analysis of these regions using JASPAR identified the high mobility group (HMG) and homeobox-type *cis*-elements near the ATAC-seq peaks (Fig. 2B, Fig. S2).

**Figure 2.**
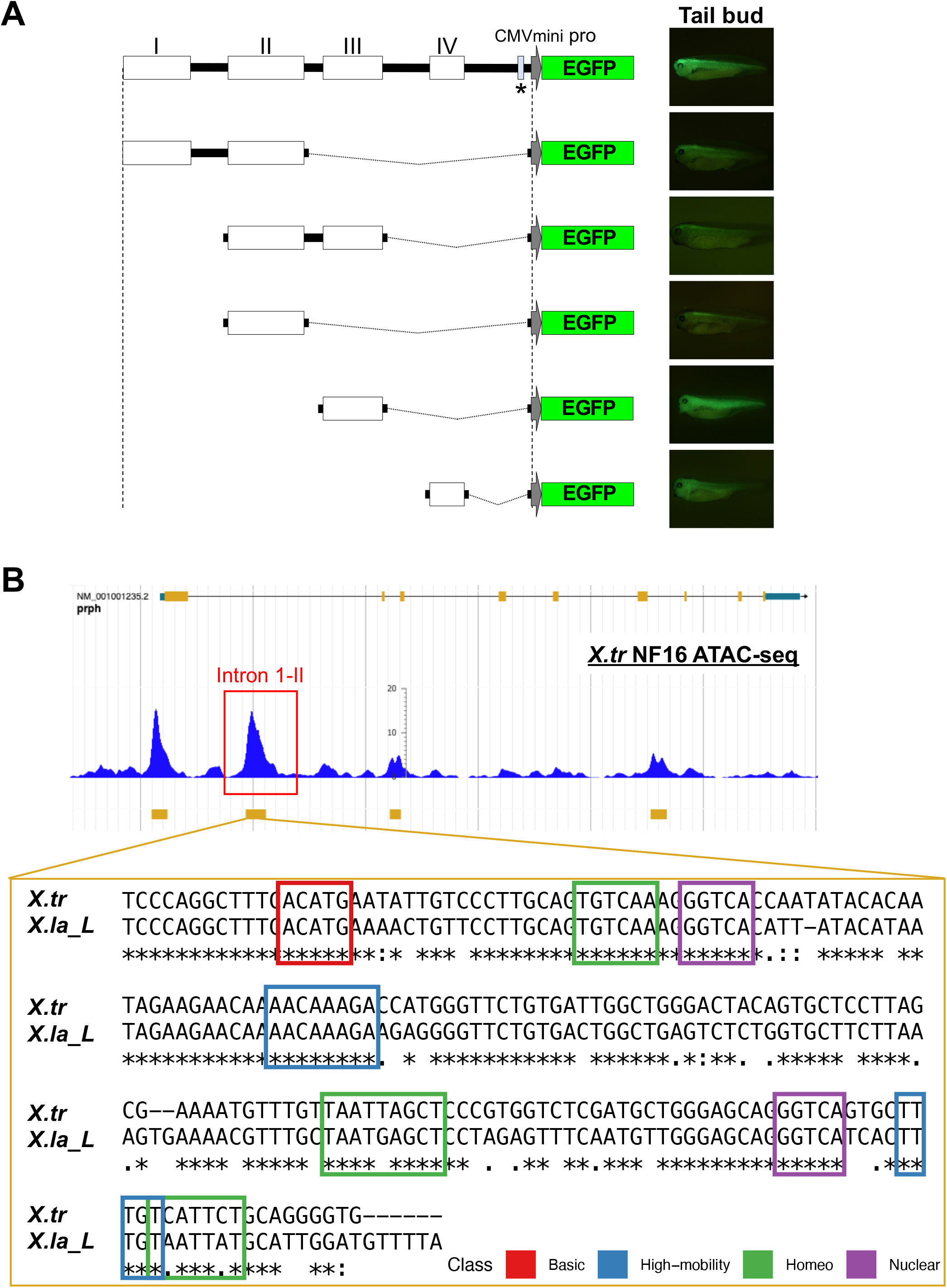
Deletion and enhancer analysis of the intron 1 of the *prph* gene in transgenic *Xenopus*. (A) Summary of sequential deletion analysis of the *prph* intron 1 sequence and representative EGFP fluorescence images of transgenic embryos. White boxes and gray arrows represent homology regions and the CMV minimal promoter, respectively. An asterisk denotes the 34 bp exon 2 sequence (light blue box). (B) *Cis*-element analysis of *X. laevis* and *X. tropicalis* homology region II (intron 1-II). Public ATAC-seq data of the *X. tropicalis prph* gene at the neurula stage revealed accessible chromatin peaks in this region, which contain putative high mobility group and homeobox-type *cis*-elements predicted by JASPAR. Abbreviations: Basic, Basic helix-loop-helix type transcription factors; High-mobility, High-mobility group domain type transcription factors; Homeo, Homeo domain type transcription factors; Nuclear, Nuclear receptors with C4 zinc finger type transcription factors.

### 2.3. *In vivo* visualization of peripheral nerves in tadpoles

We established a *prph*:EGFP transgenic line that enables *in vivo* visualization of both the CNS and PNS throughout *X. laevis* development. After hatching (st. 45), reporter expression of F1 tadpoles was clearly detected in the CNS and PNS, including the spinal cord and DRGs (arrows in Fig. 3A). Immunohistochemical analyses of tissue sections further confirmed EGFP expression in the hindbrain, spinal cord, and spinal nerves. Reporter protein was specifically detected in the ventricular layer (VL), spinal cord, and DRGs, and various neurons at the embryo and pre-metamorphic stage (st. 42 and 52 in Fig. S3 and Fig. 3B, respectively). Double immunostaining revealed co-localization of EGFP with acetylated tubulin (AcTub) in DRG, demonstrating that this transgenic line faithfully reflects neuronal distribution in both the CNS and PNS. In the PNS, EGFP signals were detected throughout the body along with peripheral nerves (Fig. 3C), allowing continuous, *in vivo* observation of innervation in living tadpoles.

**Figure 3.**
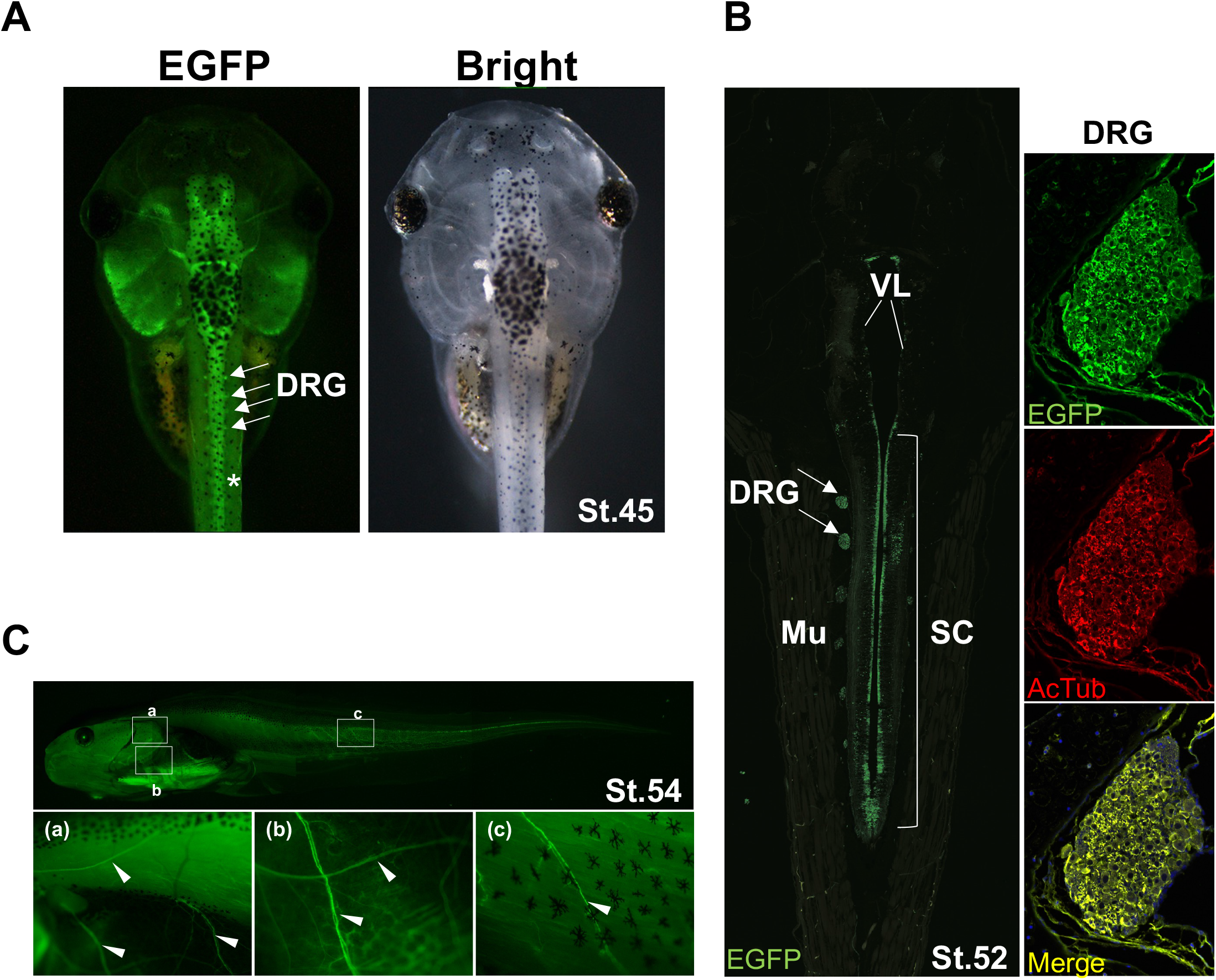
Expression pattern of *prph*:EGFP in tadpoles. *In vivo* EGFP reporter expression pattern at st. 45. Arrows indicate the dorsal root ganglia (DRGs). Immunohistochemical analysis of the reporter EGFP. At st. 52, clear expression is observed in the spinal cord (SC), ventricular layer (VL), and DRGs. A coronal section of the tail at st. 52 shows co-localization of EGFP and acetylated tubulin (AcTub) immunoreactivity. Arrows indicate the DRGs. (C) EGFP expression in the peripheral nervous system (PNS) at the pre-metamorphic stage. Upper panel: Lateral view of a st. 54 larva. Lower panel: Enlarged views of the boxed regions (a, b, c) showing peripheral nerves. Arrowheads indicate peripheral nerves. Note that an asterisk in the tail muscle (in A) marks background due to autofluorescence, where no immunosignal is observed using anti-EGFP antibody (Mu in B).

### 2.4. Innervation during limb development and regeneration

Using the *prph*:EGFP reporter line, we performed time-course analyses of peripheral innervation in tadpoles during limb development and regeneration. Sequential EGFP images of hindlimb development from st. 51 to 55 revealed the extension of peripheral nerves from the trunk DRGs into the limb bud (Fig. 4A). Initially, few nerves were found to enter into the limb bud at the onset of outgrowth; however, as patterning proceeded, extensive innervation and branching were detected distally within the developing autopod.

**Figure 4.**
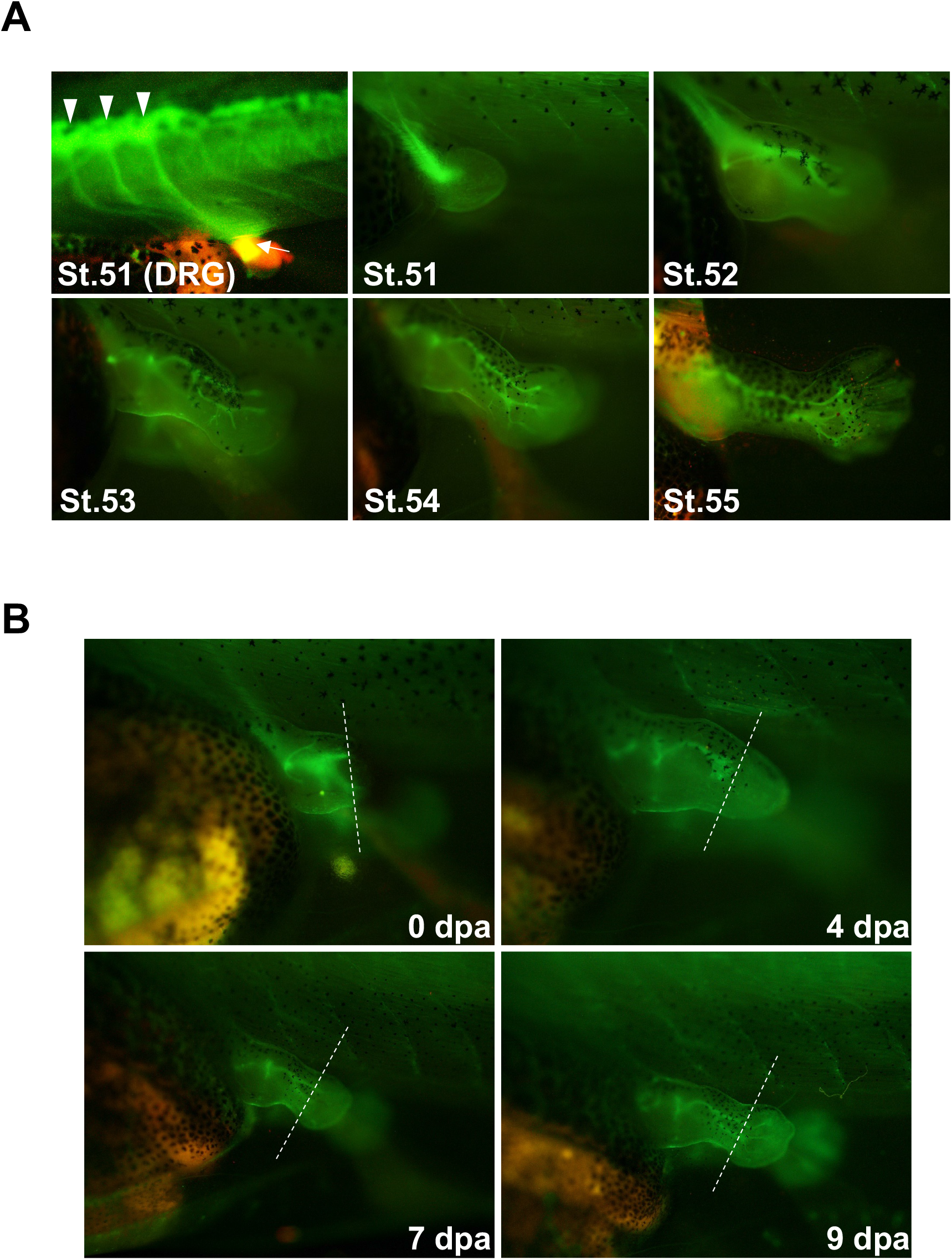
Live imaging of peripheral nerve patterns in limb development and regeneration in tadpoles. (A) Time-course imaging of *prph*:EGFP expression during hindlimb development from st. 51 to 55. At st. 51, strong EGFP fluorescence is observed in peripheral nerves extending from the dorsal root ganglia (DRGs, arrowheads) to limb bud (arrow). Subsequently, innervation follows the patterning of limb development. (B) Time-course imaging of peripheral nerves during hindlimb regeneration in tadpoles. The following time points were examined: 0 days post-amputation (dpa; immediately after amputation), 4 dpa (blastema formation), and 7–9 dpa (palette stage). Dotted lines indicate the amputation plane. Peripheral nerves do not markedly invade the blastema, with only sparse invasion in later regenerative stages.

We next observed peripheral innervation during limb regeneration. *X. laevis* tadpoles possess robust regenerative capacity and can fully regenerate amputated hindlimbs at the pre-metamorphic stage, whereas post-metamorphic froglets fail to complete regeneration, resulting in a spike structure. In larval hindlimb regeneration (Fig. 4B), the early blastema at 4 days post-amputation (dpa) exhibited markedly sparse innervation. As patterning progressed, peripheral nerve growth and branching recapitulated the patterns observed during early limb bud development. In contrast, sequential imaging of froglet limb regeneration revealed extensive invasion of peripheral nerves into the mesenchyme of the blastema during its formation (Fig. 5A). This notable innervation was confirmed by the consistent co-localization of EGFP with the axonal marker acetylated tubulin in immunostained sections of blastema (Fig. 5B). Additionally, we observed that EGFP fluorescence in the nerves was consistently stronger in the regenerating forelimb compared to the intact (non-amputated) limb of the same animal (Fig. 6A). This difference was also evident in dissected forelimb nerves from both sides (Fig. 6B), and quantitative analysis of mean fluorescence intensity confirmed a significant increase on the regenerating side (*p* < 0.01, n = 10; Fig. 6C). Mouse PRPH is predominantly synthesized in the somata and transported to peripheral nerves in an anterograde fashion (Xia et al., 2003). Consistent with this, the increased EGFP signal in the nerves likely reflects *prph* promoter/enhancer activation in the DRG during *Xenopus* limb regeneration. Therefore, these results suggest regeneration-associated activation of *prph* in peripheral nerves, possibly influenced by transcriptional changes in the DRGs.

**Figure 5.**
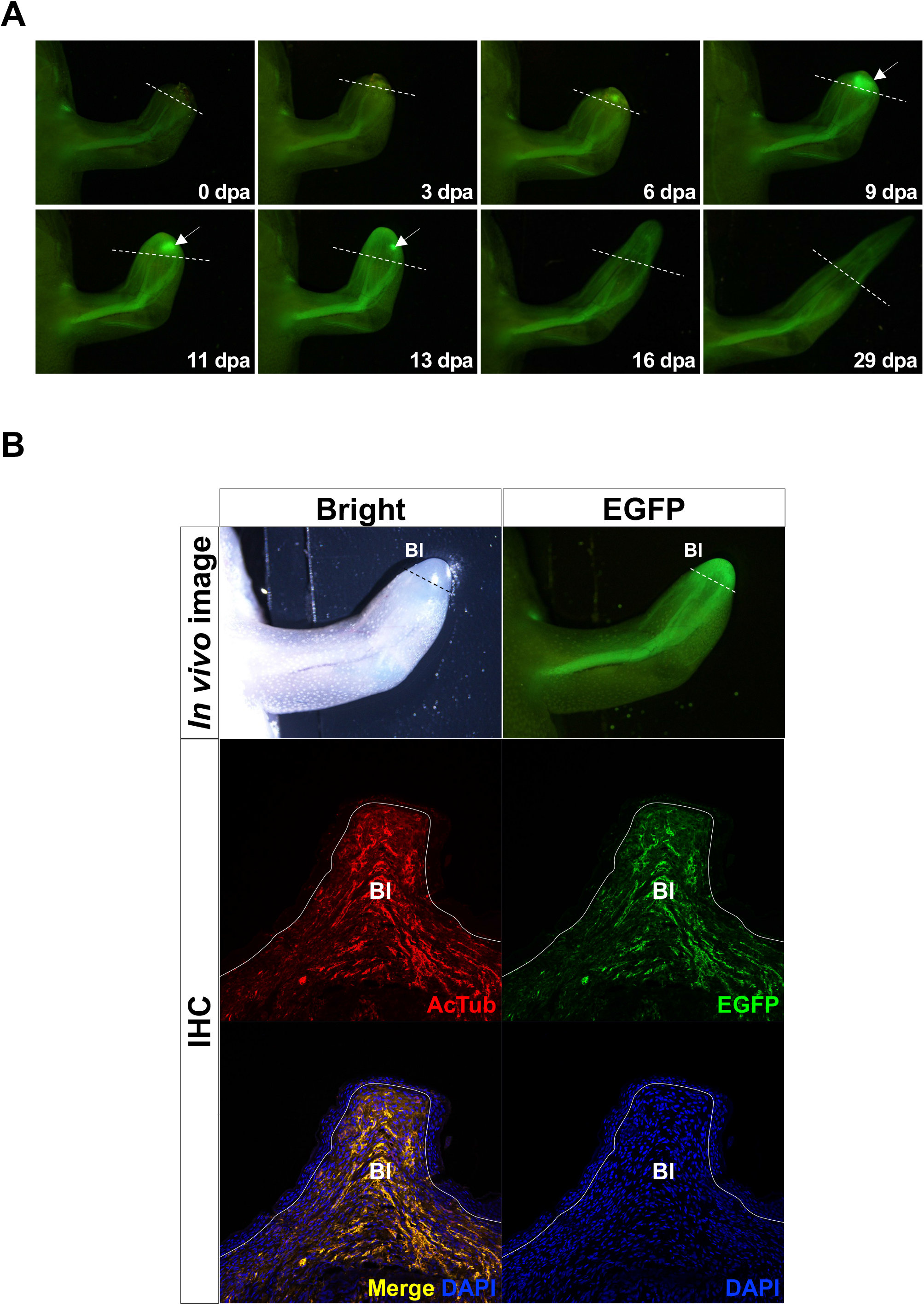
Time-course imaging of peripheral nerve patterns during forelimb regeneration in post-metamorphic froglets. (A) Time-course EGFP imaging of peripheral nerves during forelimb regeneration from 0 to 29 dpa (days post-amputation), until spike formation. Unlike larval hindlimb regeneration, extensive peripheral innervation into the blastema is observed. Dotted lines indicate the amputation plane. A bright mass of EGFP fluorescence was noted near the amputation plane (arrows in A), potentially due to excessively dense innervation. We confirmed that this signal was not observed in wild-type froglets (Figure S4), thereby ruling out the possibility of autofluorescence. This reporter fluorescence appeared transiently between 4–9 dpa and became undetectable after 11–13 dpa (Figure S5). (B) Robust peripheral innervation at the blastema formation stage (11 dpa). The upper panels show an enlarged view of *in vivo* imaging, and the lower panels show an immunostained histological section for the axonal marker acetylated tubulin (AcTub), EGFP, and DAPI. The *prph*:EGFP expression overlaps with acetylated tubulin staining, demonstrating that the reporter faithfully recapitulates PNS expression. Dotted and solid lines indicate the amputation plane and the epidermal–mesenchymal boundary, respectively. Bl, blastema.

**Figure 6.**
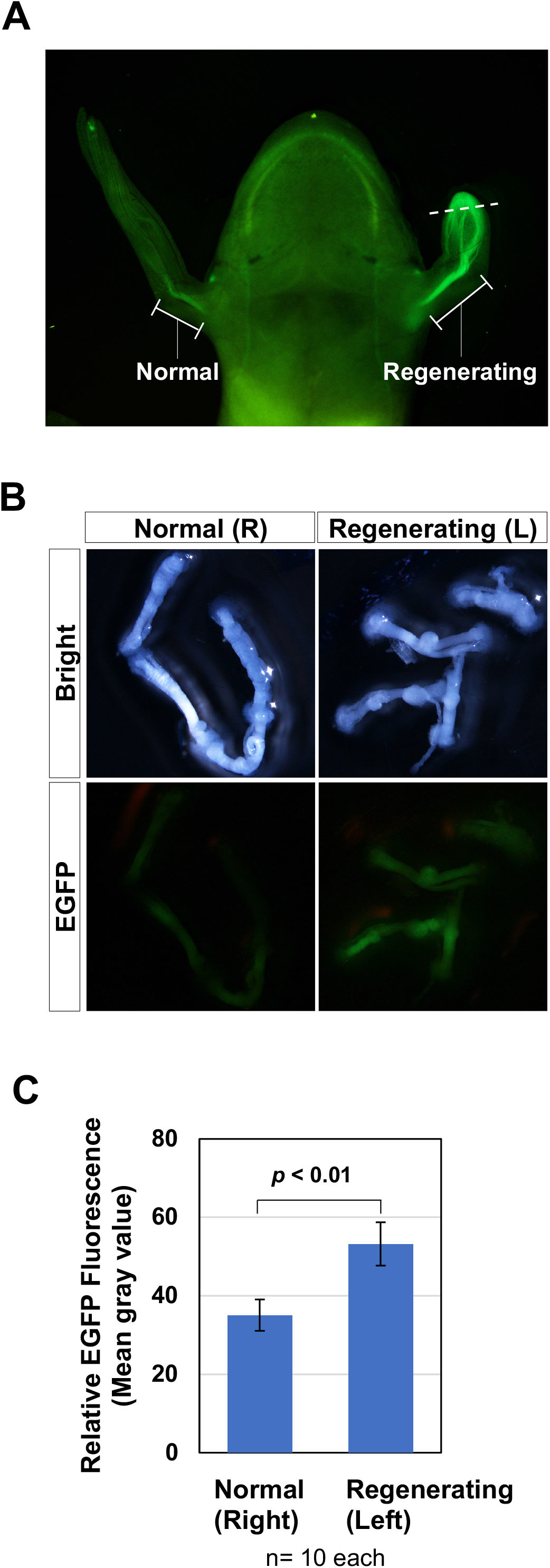
Regeneration-associated activation of *prph*:EGFP expression. (A) *In vivo* EGFP imaging in ventral view of forelimb regeneration in a froglet. The left side shows an intact right forelimb (Normal, mirroring), and the right side shows a regenerating left forelimb (Regenerating, mirroring). The EGFP signal in the nerves of the forelimb is stronger on the regenerating side. The dotted line indicates the amputation plane. (B) Dissected forelimb nerves from intact (Right, R) and regenerating (Left, L) limbs. Consistent with the *in vivo* imaging, the EGFP signal is stronger in the regenerating nerves. (C) Quantification of mean EGFP signal intensity comparing intact (Right forelimb) and regenerating (Left forelimb) nerves at 11 dpa (Mann–Whitney U-test; *p* < 0.01, n = 10 each).

## 3. Discussion

Peripherin (*PRPH*) is a type III intermediate filament protein expressed in the PNS and is implicated in critical roles during axonal development and regeneration. In this study, we established a stable reporter line in *X. laevis* that labels the PNS, enabling *in vivo* visualization of peripheral nerves during development and regeneration. The *PRPH* gene is known to respond to nerve injury (Oblinger et al., 1989; C.M. Troy et al., 1990; Wong and Oblinger, 1990) and has attracted attention for its potential role in nerve damage, including in neuropathies (Keddie et al., 2023; Romano et al., 2022). We demonstrated extensive peripheral innervation of the blastema during limb regeneration and strong activation of the *X. laevis prph* promoter/enhancer *in vivo*. Our results highlight *X. laevis prph* as a valuable marker for advancing our understanding of peripheral nerves in early development, metamorphosis and regeneration.

### 3.1 Transcriptional regulation of *prph* in development

Our reporter analysis showed that the 4.7 kb intron 1 is crucial for precise expression in the CNS and PNS across developmental stages, extending from early development to beyond metamorphosis. In particular, we found that Region II of intron 1 is essential for *prph* expression in a neuron-specific manner. In reporter transgenic mouse studies, it was also reported that intron 1 is essential for the cell type-specific expression of rat *Prph* in the DRG, enteric nervous system, and motor neurons (Uveges et al., 2002). Taken together with previous studies, our findings indicate that the *PRPH* intron 1 is evolutionarily conserved in its cell-type specificity among vertebrates. Further analysis using ATAC-seq data and JASPAR revealed neurula stage-specific chromatin accessibility peaks in the promoter region and Region II of intron 1, which contained putative HMG and homeobox-type motifs. Mammalian *PRPH* genes are strongly expressed not only in the differentiated PNS but also in the neural primordium of early vertebrate embryos, highlighting its importance during early mammalian neurogenesis. In the human *PRPH* promoter, neural crest-related transcription factor *AP-2* (*TFAP2*) motifs have been identified (Foley et al., 1994; C.M. Troy et al., 1990). HMG binding motifs were found in the promoter and intron 1 Region II, suggesting that *Xenopus prph* expression is possibly regulated by the neural crest regulator, *sox10*. Although *ebf3* is likely to directly regulate *prph* in *X. laevis* (Green and Vetter, 2011), no *EBF*-family binding motifs were detected in the regions analyzed here.

### 3.2 Differences in peripheral innervation between larval and adult limb regeneration

The *prph*:EGFP line enabled *in vivo* imaging of limb development and regeneration, revealing distinct peripheral innervation patterns between larvae and adults. In tadpoles, limb buds showed minimal innervation at early stages of development, with increases as patterning progressed; regeneration followed a similar pattern. In contrast, adult regeneration displayed extensive innervation during the blastema formation stage, potentially supporting blastema outgrowth. The importance of nerves and nerve-derived factors in limb regeneration has been extensively documented (Kumar and Brockes, 2012; Pirotte et al., 2016; Stocum, 2017). The accessory limb model (ALM) further illustrates this; ectopic blastemas can be generated by deviating the nerve into a skin wound, suggesting that the presence of the nerve is essential for blastema formation (Endo et al., 2004). Moreover, manipulations leading to hyperinnervation improve the limb regeneration capacity in *X. laevis*, producing bifurcated or multiple skeletal elements rather than a simple spike structure (Kawasumi-Kita et al., 2024; Kurabuchi, 1992; Mitogawa et al., 2018). In *X. laevis* froglets, there are nerve-dependent and - independent events during blastema formation. While the reactivation of *prx1*, a mesenchymal marker, occurs in a nerve-independent manner at the onset of limb regeneration, nerves are required for blastema outgrowth including cell proliferation, cell survival, and upregulation of *fgf8* (Suzuki et al., 2005). Supporting these findings, our *prph*:EGFP line visualized the extensive peripheral innervation into the blastema during adult limb regeneration; however, several points should be noted. First, while *Prph* expression is robust in sensory and autonomic neurons derived from the DRGs, it is reported to be limited in motor neurons in rats (Brody et al., 1989; Escurat et al., 1990). Future generation of neuronal subtype-specific lines (e.g., *hb9*/*mnx1* for motor neurons) will enable a deeper dissection of nerve dependence in regeneration. Additionally, the potential contributions of Schwann cells and glia to limb regeneration must be carefully examined using reporter lines that can label and track these cells.

### 3.3 Transcriptional regulation during nerve regeneration

In *prph*:EGFP *Xenopus*, we observed a significant increase in reporter activity exclusively on the regenerating side during froglet limb regeneration. This activation likely reflects widespread transcriptional changes in the DRG in response to axotomy. In rats, *Prph* expression is significantly upregulated in DRG neurons following sciatic nerve injury during axon regeneration (Oblinger et al., 1989; Carol M. Troy et al., 1990; Wong and Oblinger, 1990). Moreover, mammalian *PRPH* has been associated with traumatic nerve injury and neuropathies, and has been highlighted as a potential biomarker for nerve damage in the PNS (Hol and Capetanaki, 2017; Keddie et al., 2023; Manco et al., 2025; Romano et al., 2022). Our line enables *in vivo* visualization of the dynamics of peripheral nerves during axonal degeneration and regeneration, thereby providing a valuable tool for studying nerve plasticity in developmental, regenerative, and pathological contexts.

In rat PC12 cells, NGF activates *Prph* expression *via* Ets family transcription factors, which bind the distal positive element (Chang and Thompson, 1996; Thompson et al., 1992). Considering the upregulation of NGF and other neurotrophic factors (e.g., BDNF and GDNF) in the DRG after nerve injury (Lee et al., 1998; Song et al., 2008), such factors may also be induced by regeneration cues following limb amputation. Thus, we hypothesize that injury-induced neurotrophic factors upregulate *prph* expression *via* its promoter and intron 1 Region II, in concert with transcription factors such as members of the ETS family.

In this study, we focused on the *Xenopus prph* gene, which serves as a marker for peripheral nerve development and regeneration in the limb. We generated a transgenic line that allows *in vivo* visualization of *prph* expression, thereby facilitating the study of nerve development, metamorphosis, and regeneration. Peripheral nerve regeneration is highly relevant to human disorders, including traumatic nerve injury and neuropathies. Hirschsprung disease represents a neural crest-related disorder that causes defects in enteric peripheral nerves (Heuckeroth, 2018; Montalva et al., 2023). As *PRPH*, potentially regulated by neural crest transcription factors, labels enteric ganglion cells and serves as a useful diagnostic marker (Chisholm and Longacre, 2016), our transgenic line may also facilitate studies of enteric nervous system abnormalities, including those in Hirschsprung disease. Although regenerative capacity in mammals is limited, comparative studies with highly regenerative species such as fish and amphibians hold significant potential. Given that the extensive tissue remodeling during *Xenopus* metamorphosis involves a global reorganization of the peripheral nervous system (Marsh-Armstrong et al., 2004; Nakagawa et al., 2000), this transgenic line will also provide a versatile resource and tool for analyzing such large-scale peripheral nerve remodeling across development, metamorphosis, and regeneration.

## Materials and methods

### 4.1. Animals

Wild-type *X. laevis* adults were purchased from Yumesaki Youshoku or Hamamatsu Seibutsu Kyouzai, Japan. MS-222 (Sigma) was used at 0.01–0.05% for anesthesia. Tissue sampling was performed following deep anesthesia (0.1–0.2% MS-222) and proper euthanasia. Developmental stages of *X. laevis* embryos and larvae were determined according to Nieuwkoop and Faber, 1967. The animals were handled in accordance with protocols approved by the Animal Care and Use Committees of the University of Hyogo and the National Institutes of Natural Sciences. Additionally, handling was in accordance with the guidelines of Hiroshima University for the use and care of experimental animals (Former affiliation, M. S. and K.T.S.).

### 4.2. Cloning of the *X. laevis prph* gene and plasmid construction

A genomic DNA library from wild-type *X. laevis* (Suzuki et al., 2010) was screened by hybridization with a probe made from the *prph* cDNA (clone XL 79h11; XDB3). One of the positive clones was characterized by PCR and partial sequencing and was found to contain the upstream region and the first exon. The first intron sequence was amplified from the genomic DNA and cloned into the HindIII/SalI sites of the pminCMV-EGFP vector. Sequentially deleted fragments were subcloned into the same vector. A 914 bp fragment containing the first exon and 5’ UTR, and 4.7 kb fragment of the first intron were amplified and subcloned into the KpnI/BamHI and EcoRI/SalI sites, respectively, of the pEGFP(-P) vector (Suzuki et al., 2010) to generate the *prph*:EGFP construct for transgenesis. Primers used for constructing the transgenic *X. laevis*, and sequences of the promoter and intron 1 are listed in Fig. S6.

### 4.3. Transgenesis

The plasmid DNA was linearized by digestion with HindIII. Transgenic *X. laevis* were produced according to Kroll and Amaya, 1996 with modifications (Mizuno et al., 2005). Transgenic F1 offspring were obtained by natural mating or artificial insemination between F0 females and males.

### 4.4. Whole-mount *in situ* hybridization

Wild-type embryos were fixed in MEMFA solution and processed for WISH as described previously (Sugiura et al., 2009). To detect endogenous *prph* expression, EST clone XL79h11 was used as a template for the antisense probe.

### 4.5. Sectioning and immunohistochemistry

For hindbrain, spinal cord, DRG, and blastema immunostaining, tissues were collected under anesthesia and fixed with 4 % paraformaldehyde. Tissues were processed for paraffin embedding and sectioned at 8 μm. The sections were blocked with 10% fetal bovine serum in phosphate buffered saline containing 0.1% Tween20 (PBST) for 60 min at room temperature (RT) and incubated with mouse anti-acetylated tubulin antibody (Sigma, diluted 1:500) and rabbit anti-GFP antibody (Rockland, diluted 1:500) overnight at 4 °C. After washing, sections were incubated with Alexa Fluor 488- and/or 594-labeled secondary antibodies in PBST (Life Technologies, diluted 1:1000) and counterstained with DAPI or Hoechst 33342 (Life Technologies).

### 4.6. VISTA and JASPAR analysis

Comparison of *X. laevis prph* genomic sequences with *X. tropicalis* was performed using the mVISTA program (Frazer et al., 2004) (http://genome.lbl.gov/vista/) based on LAGAN multiple alignments (Brudno et al., 2003) using default parameters. For JASPAR (https://jaspar.elixir.no/) score calculation, the fragment containing the first exon to 4.7 kb of the first intron was submitted to JASPAR Scan (Rauluseviciute et al., 2024). All transcription factors in the latest version of the human CORE dataset (755 TFs in total) were used for motif analysis. The relative threshold score was set to 95 %.

### 4.7. Imaging and fluorescence intensity analysis

For *in vivo* visualization of peripheral nerves during development and regeneration, *prph*:EGFP fluorescence was observed using fluorescence stereomicroscopes (Leica MZ10F, MZ16F, or M165FC; Leica Microsystems, Germany) and digital cameras (Nikon DS-5Mc or DS-Ri1; Nikon, Japan). Fluorescence intensity was measured using ImageJ, and statistical significance was evaluated with the Mann–Whitney U-test in EZR software (Kanda, 2013).

## Supporting information

Suzuki et al_Supplementary information

## Acknowledgements

This work was financially supported by grants from the Japan Society for the Promotion of Science (JSPS), KAKENHI Grants-in-Aid for Scientific Research (B) (JP18K06257, JP21H03829, and JP25K02288 to KTS), Grants-in-Aid for Scientific Research (C) (JP18K06266 to MM); a Grant-in-Aid for Scientific Research on Innovative Areas (22124006 to TE and 25124708 to KTS), the National Institute for Basic Biology (NIBB) Japan Collaborative Research Program (25NIBB338 to MM); and the Japan Science and Technology Agency (JST), CREST program (JPMJCR 2025 to KTS). We thank Ms. Risako Kondo and Kyoka Manabe for their assistance with this research. Some experimental data were acquired in the previous affiliation, Hiroshima University.

## Declaration of generative AI and AI-assisted technologies

The authors acknowledge the use of ChatGPT and Gemini for language editing support. All AI-assisted texts were reviewed and revised by the authors. The authors take full responsibility for the contents.

