## Supplementary material for "Visualization of peripheral nerves in developing and regenerating limbs using a novel peripherin reporter line of *Xenopus laevis*": Suzuki et al_Supplementary information

**Supporting information**

**Figure S1. Deletion and enhancer analyses of intron 1 of the *prph* gene in transgenic *Xenopus***

Data from sequential deletion analyses of the *prph* intron 1 sequence are summarized in Figure 2A. The number of CNS-specific EGFP expressing transgenic embryos at the tailbud stage (EGFP positive embryos/total embryos) is indicated for each construct. White boxes, a black box, and a gray arrow represent homology regions, exon, and the CMV minimal promoter, respectively.

**Figure S2. *Cis*-element analysis of Homology Region II (Intron 1-II) of the *X. tropicalis prph* gene**

ATAC-seq peaks at the neurula stage are shown with the corresponding sequences. Predicted transcription factor binding motifs are highlighted with colored boxes. ATAC-seq data are from Xenbase. *Cis*-elements of Region II in intron 1 are predicted by JASPAR. Abbreviations: Basic, Basic helix-loop-helix type transcription factors; High-mobility, High-mobility group domain type transcription factors; Homeo, Homeo domain type transcription factors; Nuclear, Nuclear receptors with C4 zinc finger type transcription factors.

**Figure S3. Immunohistochemical analysis of EGFP reporter in the *prph* transgenic embryos.**

Transverse sections of stage 42 *prph*:EGFP embryos at hindbrain (HB) and spinal cord (SC) levels were immunostained for EGFP (green) and Islet1/2 (magenta in middle panels) or DAPI (magenta in lower panels). In the hindbrain, arrows indicate EGFP-positive trigeminal ganglion neurons. In the spinal cord, arrows and arrowheads indicate EGFP-positive dorsal root ganglia (DRGs) and EGFP-negative Rohon-Beard (RB) cells, respectively. EGFP expression is prominent in ventral neurons but absent in dorsal RB cells.

**Figure S4. No fluorescence in the wild-type froglets following amputation.** A dotted line indicates the amputation plane. Neither auto- nor nerve-specific fluorescence was observed at 10 dpa (n=3).

**Figure S5. Time-course imaging of EGFP reporter fluorescence in the *prph* transgenic froglets following amputation.** Dotted lines indicate the amputation plane. Arrows denote transient and ectopic EGFP expression. These ectopic signals appeared 4–9 dpa, then diminished and became undetectable by 13 dpa (total n = 6/6 including Figure 5A).

**Figure S6. Sequences of the *X. laevis prph* promoter, intron 1, primers, and vectors used for construct generation**

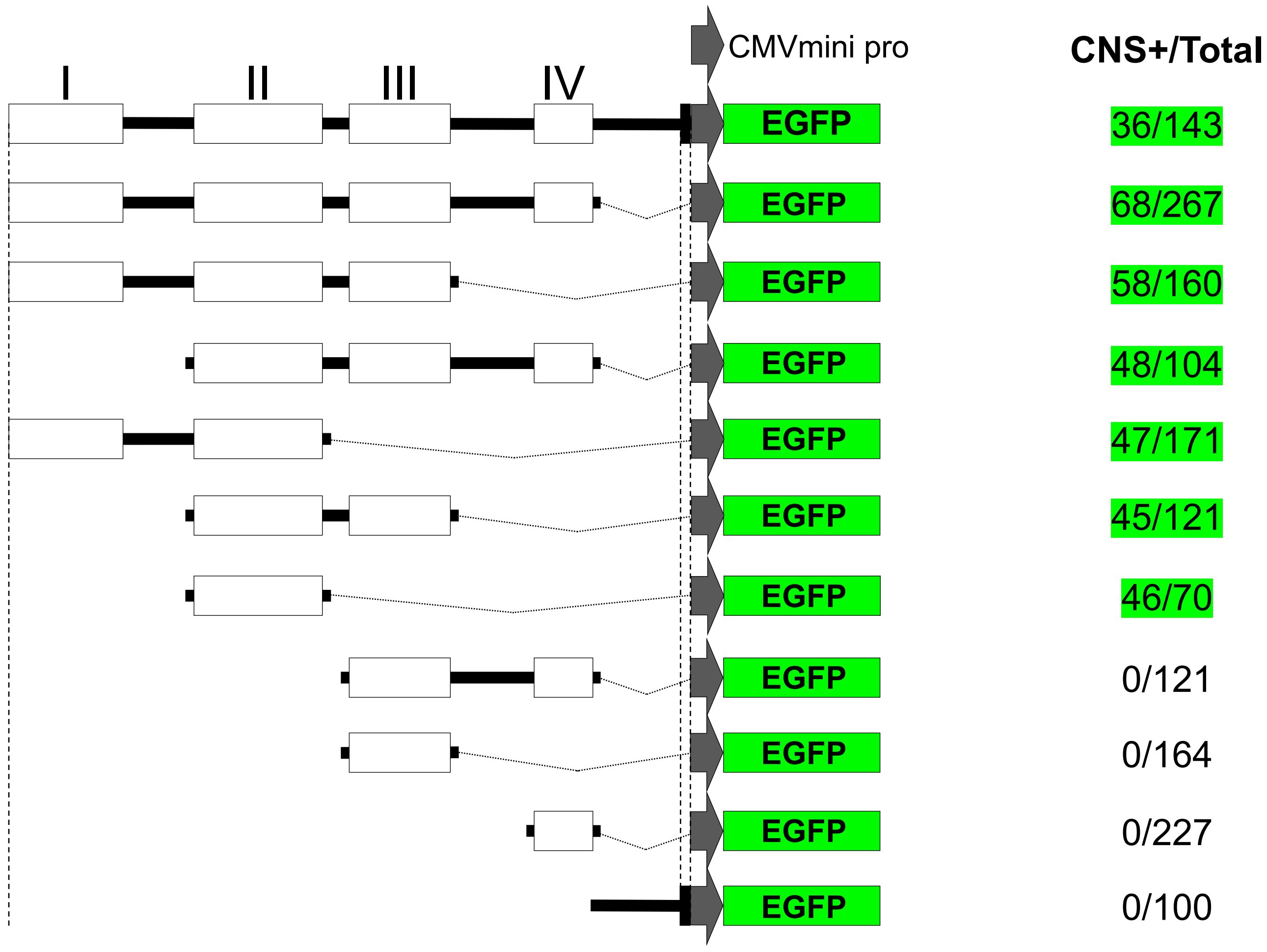
CMVmini pro
CNS+/Total
I
II
III
IV
EGFP
36/143
68/267
58/160
48/104
47/171
45/121
46/70
0/121
0/164
0/227
0/100

**Figure. S1**

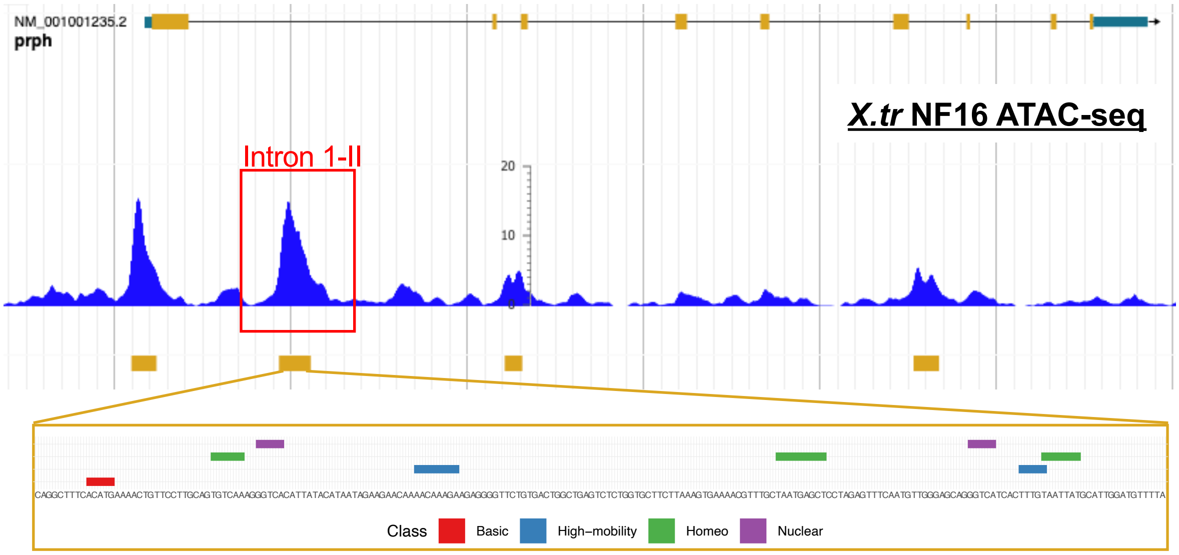


| motif | TF | score | motif | TF | score |
|---|---|---|---|---|---|
| 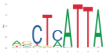 | POU6F2 | 14.799452 | 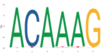 | SOX10 | 11.610084 |
| 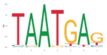 | POU6F1 | 12.783767 | 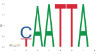 | DLX5 | 11.605389 |
| 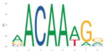 | SOX4 | 12.535857 | 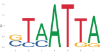 | HOXA3 | 11.563704 |
| 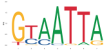 | HOXB7 | 12.223566 | 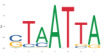 | MEOX1 | 11.529902 |
| 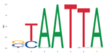 | LHX5 | 12.081846 | 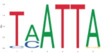 | HOXA2 | 11.304736 |
| 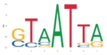 | HOXB1 | 11.695127 | 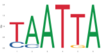 | EMX1 | 11.233512 |
| 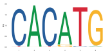 | MXI1 | 11.675601 | 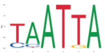 | EMX2 | 11.207822 |

**Figure. S2**

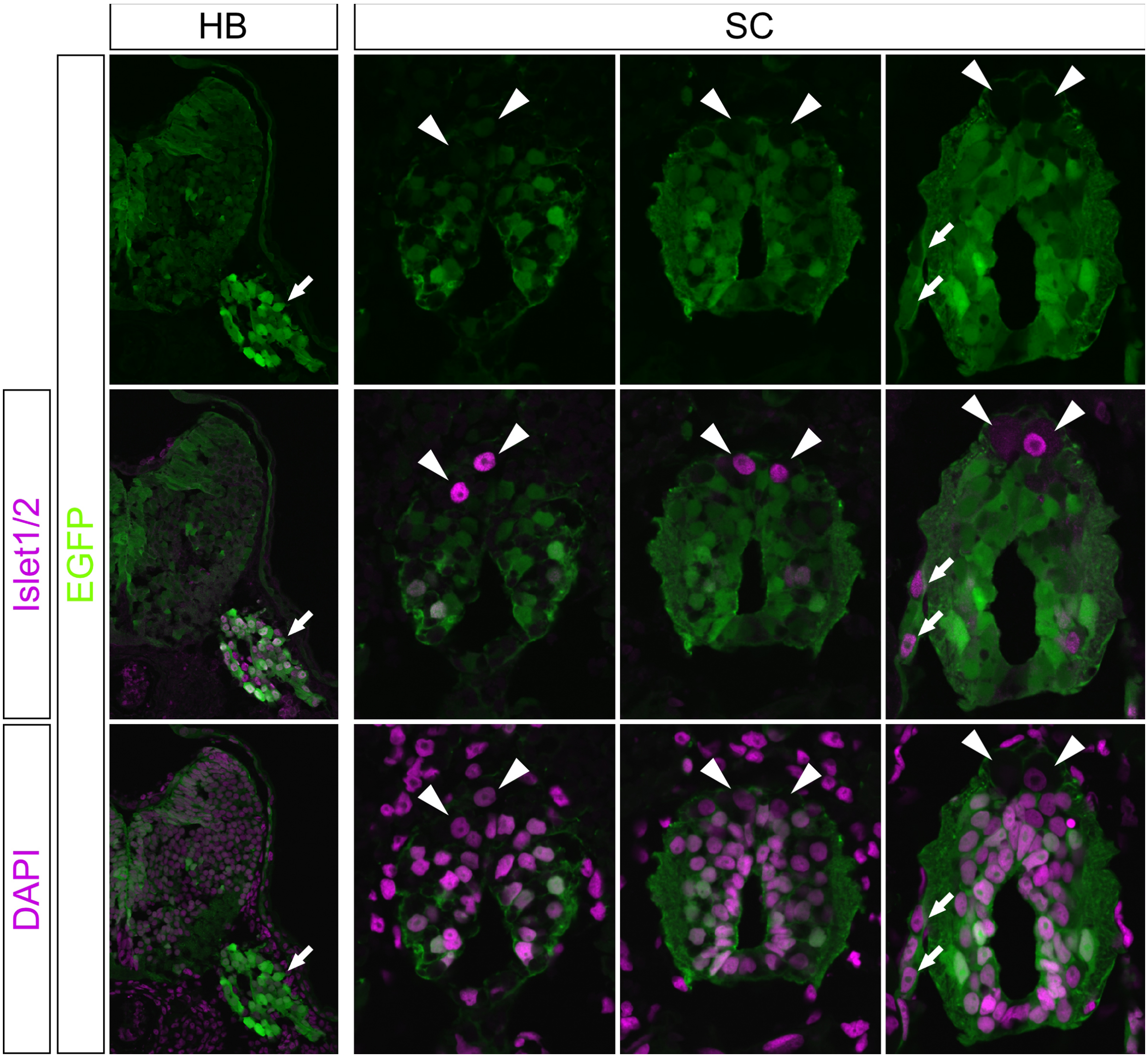


**Figure. S3**

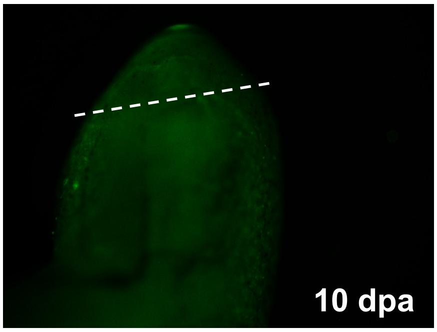


Figure. S4

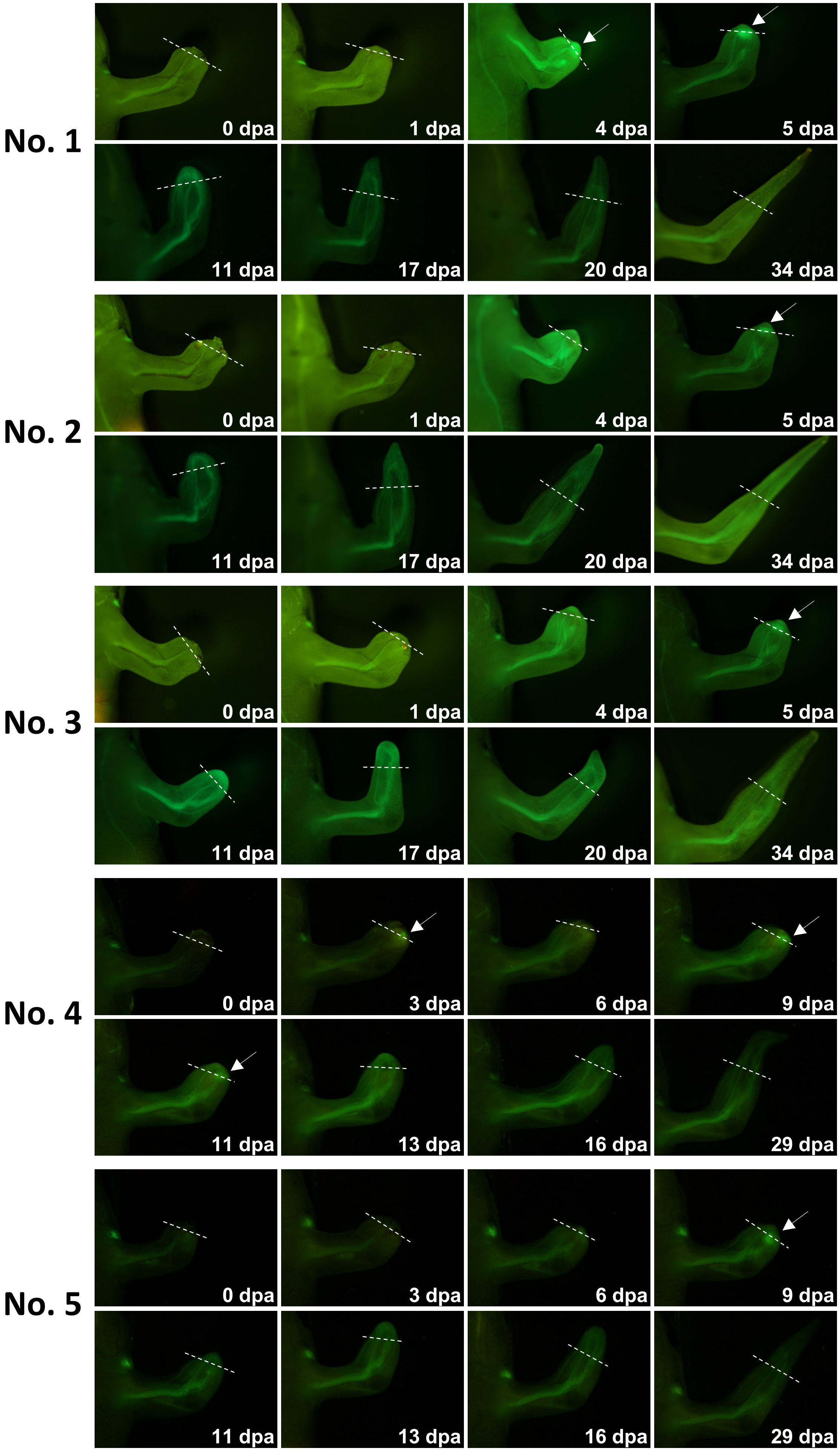
No. 1
0 dpa
1 dpa
4 dpa
5 dpa
11 dpa
17 dpa
20 dpa
34 dpa
No. 2
0 dpa
1 dpa
4 dpa
5 dpa
11 dpa
17 dpa
20 dpa
34 dpa
No. 3
0 dpa
1 dpa
4 dpa
5 dpa
11 dpa
17 dpa
20 dpa
34 dpa
No. 4
0 dpa
3 dpa
6 dpa
9 dpa
11 dpa
13 dpa
16 dpa
29 dpa
No. 5
0 dpa
3 dpa
6 dpa
9 dpa
11 dpa
13 dpa
16 dpa
29 dpa

Figure. S5

### Sequence of 914 bp *prph.L* promoter

gtcgacggatcgcggccgcgaattctcgatcatctctgcagggggctaggggaataacaattaggggagtcaaccctgcaaatactggtggtctaggagaagggggagctgccatgtgca
atttatgtgagattatttatttaaaaaaacttgatttaatagggtcgtgaaaaaactcgataaaatcaagctaaaacaaattgtacaattctgaaaaaagtcagattttccggctaaacc
cagcgcagaccacgataacttgaaattaggatagctgcctctcccattgacaggtctgagatggcagatttttggattctatctttttgcagaattgagtataataaatctcgaaaaaaa
tctagtttttttttcctctaaaaattagagttttctagtgataaaaaaatgcaaatattaagagatttaacaactcgtgctttaataaataacccccaacatctctggtgttatggggcc
ctaaaaatattttgctgtgcggtcccttcattttagttacaacattaaaggggatggacaggggacatctaaatcctatattcatgtgaatataaatatattgttagttgtttctgagct
gtgttccattcacttacagccacagagcataagccgttgaatagaaggatgaaagtaaatctgagtgctgtaggaagatgcacagagtgagttcagcatataatgggtgggtgtggtgga
tgcagatcccccctcctccctctgctcagggttatgcatctatgtggttccacagggaggggctgagtctgcacaataagatgctgcatgctggcttataaagcagcctgccatactctca
gactttcttctcaacatcactcgagaaCTAAGTACTAACATCCAGCTTAAGTGCTTGGGAATTCCTTGCTCAT

Blue, 5´ UTR

**Figure. S6**

### Sequence of *prph.L* Intron I (4,718 bp)

gtaaggtgccagggggtttagaggcagatggaggaatgtagttatttctgagggggttataagtatacaaacatgtgaagaacttctaagaagtctgaaacatttcattggtgatggggt
P1F ->
ggccatatagatatacaaaagagaagaaactgaccagtgcatattgcctttccccatattttatttgtaattaaatttttgcatatattttcaaaagcagttttaggtaaagttcatata
gtacccctattaacctctggttgtatgtttataggctggtattccttgttaggtggcatttcgtgggaaaagtgtctcctggccaaagtattggtcacacagactgattctccctgactg
tagaatttatatgggagagatctcaagaaccaggcctatttctgaaaacataagacccctccatgggtgggtgtctgttatgtgtcatgtggtggcattagattagatgcgggcagtcta
gacataggcagggtgtggccatgccactaaatattatgctctgtatcttctgtggcttactttgctacttctactgttcatcctggaacaatgctgataaagggaatgtactgaaggttt
taacaggcaacttttataatacaaaatattactgccaaagaaataactgttacactatgccatgtaactccttgttaacatgtattttatttgtatataatacacaaaagtcatgaata
<- P1R
tcttgtaaattatatccttataaacggtgagtagtgatgtcatcagttataaacggtgagtagtgatgtcatttctgtcacatgactcactaaaatttgtgtattataataaagtacccc
cagttgtaaaatatgaggatattataagttacctcggagttccatgaccatataaaagcacgaggcctcgtgtttttatatggtcatgaaactcctcggtaacttataatatccttatat
tttacaagaggggtactttattcactatacaacaaatgatccagtccagtcttgttttcttataacagtggcggaagagggaaatacttataatgaagagatggtatcaacaacagccc
ttgctgcattactattctcagcttaaaataagtaatgccacatgttatagttcagcaacaaaaagtacttgttctattacaatgtatatccatgtgataatgaacaaaaacgataaattc
P2F ->
agaaggttgatttcattattccaatcagactagactggttagttctagcttcaaattagttgtcaagggttacaggatcaaggcaaactttatggccgtttgtgcaatacgcccttgctt
tcccaggctttcacatgaaaactgttccttgcagtgtcaaagggtcacattatacataatagaagaacaaaacaaagaagaggggttctgtgactggctgagtctctggtgcttcttaaa
gtgaaaacgtttgctaatgagctcctagagtttcaatgttgggagcagggtcatcactttgtaattatgcattggatgttttatttttttactaacaaatgcacagaaagccagtttcctc
tgtgacaatgtggttgcagttgcaccaatgtgtgatctcattgatcctttgctaatcattggttaatccttcccccctagtgctcttatcttacataggtcattgcacagctggccactct
cttacccactccctatgggcaagattaaaggttaggggaagagtgcagcaccagtatgggcagaagttgaaaagcacagatgtatggataatggcaaaaagaacaatgaataaaggtggg
<- P2R
aatgggaatacaggccgtgagcactgcatcaatgagctgatcctcattttaacggaatttttaaacttgtccgattaatatctggctgattttcggccagttattaatcaggtagaccca
tcagcgggccctatagacaggcaggtaagctgttgactctgagcctgagtccaaatctgcctgtgatttcaggaatgctaatcacactgtgttactactaggaaatgaaacattacaatc
P3F ->
ccttagtttttcagagcatcattttgtttcaaaattcataattggacagtaccataatcctttaggctgctgccatacaggtcatattccccccctgccattatttagccatctgagtaac
acccaagctgcctgttattggctttaattttggccccaaaagacagatgtaagcatttttaggctacatttagtgacaggtgtttttcactatctccagtaacaggtggcacacccctgt
gccatcattctgaagtgcaactaacttctgtttcttttcactttggtataatggaaattataggttgttggctccataaatgtttttgttaaaggcctttaaaaatagtatgtattgtat
tcacatttatagtaaaatgtcaaatagttccatgtggaaaaaaagaagaataacttatatttagagaattgtcatttaggagattgggtgtgtgagcatagacacttattgttctggtac
gcaagactgggtgtgtctatggggggcaggatgaatggtaaaatatctgcaacagggctggcacaatctgcccaagatgatattgagtgttcatatgagtgttggtttatgtatagataac
tggataaacgtgtccaagtcgcaaataattactctaggctgctgaagcatttataagaaacaaatgctttggaaacaacttttactggccaaactagtgctgttcctcttgatgccactg
<- P3R
ttgtttgcagggataaaaaggtaggtagaaaggttagcatctgtgacagagagtgtttatggactgtctggtggctgatagatgcagtcatccaaatcaggcttagtcagataacattca
tggtggggcagtgtacggccatcttaactgtctgattccagcttgagacaacagattgcaatacaacagaaagtgtgattcagtggtgtaactattcagagccgtccctccctagcaaaa
aacaaatctttaggggtgtcccctacctgcctgcataccctgctcctgcctacaggtaggagagcagcaaaagagggggggatttctttgcaactatagaggacgcagaccctgtggtctg
cgcccttatatctagttctacttatatccagactgtgttagaaaaagtcaaacctattcatggaaaataattattcaacatcttttacaatatggttgcatgtctacagcaattgctgtt
P4F ->
ccacttaatccacatatacaatacacccttggggatttatagagctcagaatatgagcaatactaaaatctatgggcaggttgtgaaaaaacccataacttctatacattctgttctctg
cgtactctactaaaactgtacatacaacacagtccaacattatgacttgtccactccagataaactttgtagctgattgccccatttaaggggctaaagtgactgcttgccagatatctt
gccaggaattggctagatatctctcggggaggcgggggtagttaaacatttgaatgctgcattgggctactgatttggtcctttgctaccagatcaccataggtctctgttatggtctg
<- P4R P5F ->
cacgttgtattggaggctaaacaatcagatcagcttgactcagggtgactgaatcaagcaaagagtcacttgtttgtctaacttgcttaaaggaaaactataccccccaaaatgaatactt
aagcaacagatagtttatatcaaattgaatgacatattaaagaatcttaccaaactggaatatatatttacataaatattgcccttttacatctcttgccttgaaccaccatttcgtgac
tctatctgtgctgcctcagagatcacctgaccagaaatactacaacactaactgtaacaggaagaagtgaggaagcaaaaggcagaactctgtctgttaattggctcatgtgaccttaca
tgtggtttgtatgtgtgcacagtgaatcttacgatcccaggggggcggcccttattttttaaaatggcaattttctatttatgattacccaatggcacatactactaaaaaagtatattat
tatgaaaatggttcatttacatgaagcagggttttacacatgagctgttttactcagtatcttttaatagagacctacattgtttggggggtatagttttcctttaataagataatctta
aaacaggtaaatgaccttccttttttgcttcttttaattcatattaaaaaaggtgatgcaaagctaattaaatgaccttcctttttgcttctgtaattcaaagcctcaaaacaacctcaga
tgaacgtgaaattggttagagtgcctttctaaaaacctttgcaatttacattatttttcaagatcttgacagtggctggactgtacaaatggtttaaatgggggtacatagctgaaatga
ctgtacaagaaatgggatgacatggctgaactggagtggtgtgcaaaatatattccaaatttacttctaccatattgagatttggaatgcagtgtagcctgaagaaagtggagagggaag
gttaatgccttattttatcgattacataaaaagagctcacaagtggtaatctttagacttggacaccttatggtatttgttaccatggcttttagtgatttttgtttcttttattcctgg
ccagGCTAGATGAAGAAGTTCACAAACGGGAAGATGCAGAGAATAATCTAGTCCTGTTTAGAAAGgtgagtcaaaaaaaaagagtaagaaacacctctgtctgatccagtcaggttaccat
<- Ex2R

Primer sequences are underlined.

**Figure. S6**

### Primers for subcloning of the first intron fragments

| Fragment | Forward primers | Reverse primer |
|---|---|---|
| I II III IV | P1F | P4R |
| I II III | P1F | P3R |
| II III IV | P2F | P4R |
| I II | P1F | P2R |
| II III | P2F | P3R |
| III IV | P3F | P4R |
| II | P2F | P2R |
| III | P3F | P3R |
| IV | P4F | P4R |
| V | P5F | Ex2R |

| | Primer list | |
|---|---|---|
| Name | sequence | Tm |
| peripherinG1-exon2-Rv (Ex2R) | 5-TGC ATC TTC CCG TTT GTG AAC TTC TTC ATC-3 | 66 |
| PRPHint1P1-Fw-Hind3 (P1F) | 5-GCG AAG CTT AAG GTG CCA GGG GGT TTA GAG-3 | 69 |
| PRPHint1P2-Fw-Hind3 (P2F) | 5-GCG AAG CTT GCC ACA GGT TAA AGT TCA GCA-3 | 69 |
| PRPHint1P3-Fw-Hind3 (P3F) | 5-GCG AAG CTT GGC AGG TAA GCT GTT GAC TCT-3 | 68 |
| PRPHint1P4-Fw-Hind3 (P4F) | 5-GCG AAG CTT CCA GAC TGT GTT AGA AAA AGT-3 | 62 |
| PRPHint1P5-Fw-Hind3 (P5F) | 5-GCG AAG CTT GAA TGC TGC ATT GGG CTA CTG-3 | 71 |
| PRPHint1P2-Rv-Sal1 (P2R) | 5-GCG CGT CGA CCC ATT CCC ACC TTT ATT CAT-3 | 72 |
| PRPHint1P3-Rv-Sal1 (P3R) | 5-GCG CGT CGA CTT GGA CAC GTC TAT CCA GTT-3 | 70 |
| PRPHint1P4-Rv-Sal1 (P4R) | 5-GCG CGT CGA CGT TTA ACT ACC CCC CGC CTC-3 | 75 |

**Figure. S6**

### Partial sequence of pEGFP(-P)

```
                                                 45 EcoRI  55 SalI
         10 Eco47III                38 HindIII  50 PstI     61 KpnI         76 BamHI
         |                          |      |    |    |      |              |
TAGTTATTTAGCGCTACCGGACTCAGATCTCGAGCTCAAGCTTCGAATTCTGCAGTCGACGGTACCGCGGGCCCGGGATCCACCGGCCGGTCGCCACCATGGTGAGCAAGGGCGAGGAGC

TGTTCACCGGGGTGGTGCCCATCCTGGTCGAGCTGGACGGCGACGTAAACGGCCACAAGTTCAGCGTGTCCGGCGAGGGCGAGGGCGATGCCACCTACGGCAAGCTGACCCTGAAGTTCA

TCTGCACCACCGGCAAGCTGCCCGTGCCCTGGCCCACCCTCGTGACCACCCTGACCTACGGCGTGCAGTGCTTCAGCCGCTACCCCGACCACATGAAGCAGCACGACTTCTTCAAGTCCG

CCATGCCCGAAGGCTACGTCCAGGAGCGCACCATCTTCTTCAAGGACGACGGCAACTACAAGACCCGCGCCGAGGTGAAGTTCGAGGGCGACACCCTGGTGAACCGCATCGAGCTGAAGG

GCATCGACTTCAAGGAGGACGGCAACATCCTGGGGCACAAGCTGGAGTACAACTACAACAGCCACAACGTCTATATCATGGCCGACAAGCAGAAGAACGGCATCAAGGTGAACTTCAAGA

TCCGCCACAACATCGAGGACGGCAGCGTGCAGCTCGCCGACCACTACCAGCAGAACACCCCCATCGGCGACGGCCCCGTGCTGCTGCCCGACAACCACTACCTGAGCACCCAGTCCGCCC

TGAGCAAAGACCCCAACGAGAAGCGCGATCACATGGTCCTGCTGGAGTTCGTGACCGCCGCCGGGATCACTCTCGGCATGGACGAGCTGTACAAGTAAAGCGGCCGCGACTCTAGATCAT
```

Green, EGFP coding sequence

### Partial sequence of pminCMV-EGFP

```
                                                        56 SalI
          11 Eco47III                39 HindIII  51 PstI     62 KpnI
          |                          |           |    |      |
TAGTTATTCTAGCGCTACCGGACTCAGATCTCGAGCTCAAGCTTCGAATTCTGCAGTCGACGGTACCGCGGGCCCGGGTCGAGGTAGGCGTGTACGGTGGGAGGCCTATATAAGCAGAGC

                                                                                                                  235 BamHI
                                                                                                          227 KpnI
                                                                                                          |       |
TCGTTTAGTGAACCGTCAGATCGCCTGGAGACGCCATCCACGCTGTTTTGACCTCCATAGAAGACACCGGGACCGATCCAGCCTCCGCGGCCCCGAATTCGAGCTCGGTACCCGGGATCC

ACCGGCCGGTCGCCACCATGGTGAGCAAGGGCGAGGAGCTGTTCACCGGGGTGGTGCCCATCCTGGTCGAGCTGGACGGCGACGTAAACGGCCACAAGTTCAGCGTGTCCGGCGAGGGCG

AGGGCGATGCCACCTACGGCAAGCTGACCCTGAAGTTCATCTGCACCACCGGCAAGCTGCCCGTGCCCTGGCCCACCCTCGTGACCACCCTGACCTACGGCGTGCAGTGCTTCAGCCGCT

ACCCCGACCACATGAAGCAGCACGACTTCTTCAAGTCCGCCATGCCCGAAGGCTACGTCCAGGAGCGCACCATCTTCTTCAAGGACGACGGCAACTACAAGACCCGCGCCGAGGTGAAGT

TCGAGGGCGACACCCTGGTGAACCGCATCGAGCTGAAGGGCATCGACTTCAAGGAGGACGGCAACATCCTGGGGCACAAGCTGGAGTACAACTACAACAGCCACAACGTCTATATCATGG

CCGACAAGCAGAAGAACGGCATCAAGGTGAACTTCAAGATCCGCCACAACATCGAGGACGGCAGCGTGCAGCTCGCCGACCACTACCAGCAGAACACCCCCATCGGCGACGGCCCCGTGC

TGCTGCCCGACAACCACTACCTGAGCACCCAGTCCGCCCTGAGCAAAGACCCCAACGAGAAGCGCGATCACATGGTCCTGCTGGAGTTCGTGACCGCCGCCGGGATCACTCTCGGCATGG

ACGAGCTGTACAAGTAAAGCGGCCGCGACTCTAGATCATAATCAGCCATACCACATTTGTAGAGGTTTTACTTGCTTTAAAAAACCTCCCACACCTCCCCCTGAACCTGAAACATAAAAT
```

Underline, CMV minimal promoter
Green, EGFP coding sequence

**Figure. S6**
